# Foxq2 determines blue cone identity in zebrafish

**DOI:** 10.1101/2021.03.14.435350

**Authors:** Yohey Ogawa, Tomoya Shiraki, Yoshitaka Fukada, Daisuke Kojima

## Abstract

Most vertebrate lineages retain a tetrachromatic visual system, which is supported by a functional combination of spectrally distinct multiple cone photoreceptors, UV, blue, green, and red cones. The blue cone identity is ensured by selective expression of blue (*sws2*) opsin, and the mechanism is poorly understood because *SWS2* gene has been lost in mammalian species such as mouse, whose visual system has been extensively studied. Here we pursued loss-of-function studies on transcription factors expressed predominantly in zebrafish cone photoreceptors and identified Foxq2 as a core factor driving *sws2* gene expression. *foxq2* is expressed only in the blue cone, and loss of *foxq2* selectively abolishes *sws2* expression. Comparative genomic analysis revealed that a wide range of vertebrate species retain both *FOXQ2* and *SWS2* genes. We propose that FOXQ2-dependent *SWS2* expression is a prevalent regulatory mechanism that was acquired at the early stage of vertebrate evolution

## Introduction

Most vertebrates have highly developed camera-type eyes with duplex retinas equipped with rod and cone photoreceptor cells (*1–3*). Rods with a higher light-sensitivity respond to single photons and mediate scotopic vision under twilight conditions at night. In contrast, cones show a relatively lower sensitivity without saturating in brighter light and mediate photopic vision under a daylight condition. Color discrimination is established by a combination of spectrally distinct cone subtypes, each expressing a single cone opsin out of four subfamilies: UV- (SWS1, wavelength of maximum sensitivity [λ_max_]: 360-420 nm), blue- (SWS2, λ_max_: 400-470 nm), green- (RH2, λ_max_: 460-510 nm) and red-sensitive opsins (LWS, λ_max_: 510-560 nm) (*4, 5*). Most vertebrates retain the tetrachromatic visual system organized by the four cone opsin subfamilies. A full set of genes encoding the four cone opsins are present in the southern hemisphere lamprey, a jawless vertebrate belonging to the earliest-branching vertebrate group (*6*). This fact supports the idea that the last common ancestor of vertebrates should have possessed color vision based on the four cone opsin subfamilies (*4, 7*).

Retinal progenitor cells differentiate into all types of retinal neurons in a temporal order, conserved among many species (*8–10*). In the later process, transcription factors regulate photoreceptor-specific gene expression. Cone-rod homeobox (Crx) is an upstream transcriptional regulator for both rod and cone photoreceptors (*11, 12*). A rod master regulator, neural retina leucine zipper (Nrl), and its downstream factor, Nuclear Receptor 2E3 (Nr2e3), enhance rod-specific gene expression and repress SWS1 expression (*13, 14*). With regard to cone subtypes, thyroid hormone receptor beta (Thrb) is a master transcriptional regulator for expression of LWS opsin and responsible for differential expression between LWS and SWS1 opsins in mice (*15*), zebrafish (*16*), and human (*17*). Another transcription factor, T-box 2b (Tbx2b), plays an essential role in SWS1 opsin expression in zebrafish (*18*). On the other hand, much less is known about a regulatory network governing expression of the middle wavelength-sensitive opsin genes, *sws2* and *rh2*, which have been lost in most mammalian species.

In zebrafish, a tetrachromatic freshwater fish, we found that *sine oculis* homeobox 7 (Six7) is required for expression of all the four subclasses of *rh2* genes (*rh2-1*, *rh2-2*, *rh2-3* and *rh2-4*) (*19, 20*), which are tandemly arrayed, expressed in different cone cells, and spectrally distinct from each other (*21, 22*). Six7 and its homolog Six6b control *sws2* expression as well (*23*). The cone-enriched transcription factors, Six7 and Six6b, share common DNA-binding sites in both *rh2* and *sws2* gene loci, and Six6b overexpression rescues the reduced level of *rh2* expression in *six7* deficient fish (*23*). The overlapping functions between Six6b and Six7 for *sws2* and *rh2* expression imply that an additional factor(s) directs selective expression of either *sws2* or *rh2* gene and determines the identities of the middle wavelength-sensitive cone subtypes. In the present study, gene expression profiling with isolated rods and cones enabled us to identify a list of cone-enriched transcription factors. Our *in vivo* functional analyses revealed a core transcriptional network, in which Foxq2 acts as a downstream regulator of Six7 and regulates *sws2* expression. We demonstrate that Foxq2 is a terminal selector determining SWS2 cone identity during development of the middle wavelength-sensitive cone subtypes.

## Results

### A severe reduction of *sws2* expression in *foxq2* mutant zebrafish

Six6b and Six7 are predominantly expressed in zebrafish cone photoreceptors and responsible for expression of the middle wavelength-sensitive opsin genes, *sws2* (blue) and *rh2* (green) (*23*). To identify a transcription factor(s) that governs differentiation between SWS2 and RH2 cone subtypes, we searched for cone-specific genes by comparing gene expression profiles between cones and rods. These photoreceptor cells were purified from the retinas of transgenic adult zebrafish, each of which express EGFP in all cone subtypes [*Tg(gnat2:egfp)*] or in rods [*Tg(rho:egfp)*] (*19*). The cone- or rod-enrichment in these purified samples was validated by RT-qPCR analyses of cone- and rod-specific transducin alpha-subunit genes, *i.e.*, *gnat2* and *gnat1*, respectively (Fig. 1A). A subsequent microarray analysis revealed approximately 500 genes showing more than 10-fold higher expression in cones than in rods (Data S1). These cone-enriched genes included four transcription factors, *foxq2*, E2F transcription factor 7 (*e2f7*), nuclear factor 1A (*nfia*), and nuclear receptor 2F6b (*nr2f6b*), the roles of which in photoreceptor development were not known. Genomic loci of these genes harbor Six6b- and Six7-binding sites as revealed by our ChIP-seq analysis (*23*) (Fig. S1), implying that some of these cone-enriched transcription factors mediate a regulatory function(s) downstream of Six6b and Six7. In addition, we paid attention to two cone-enriched transcription factors, *tbx2b* and *thrb.* They are known to be required for expression of *sws1* (*18*) and *lws* (*16*), respectively, but their contributions to the *sws2* and *rh2* gene expression have not been well characterized. Cone-enriched expression of these six transcription factors was verified by RT-qPCR analyses with purified cone and rod cells (Fig. 1A).

**Fig. 1.**
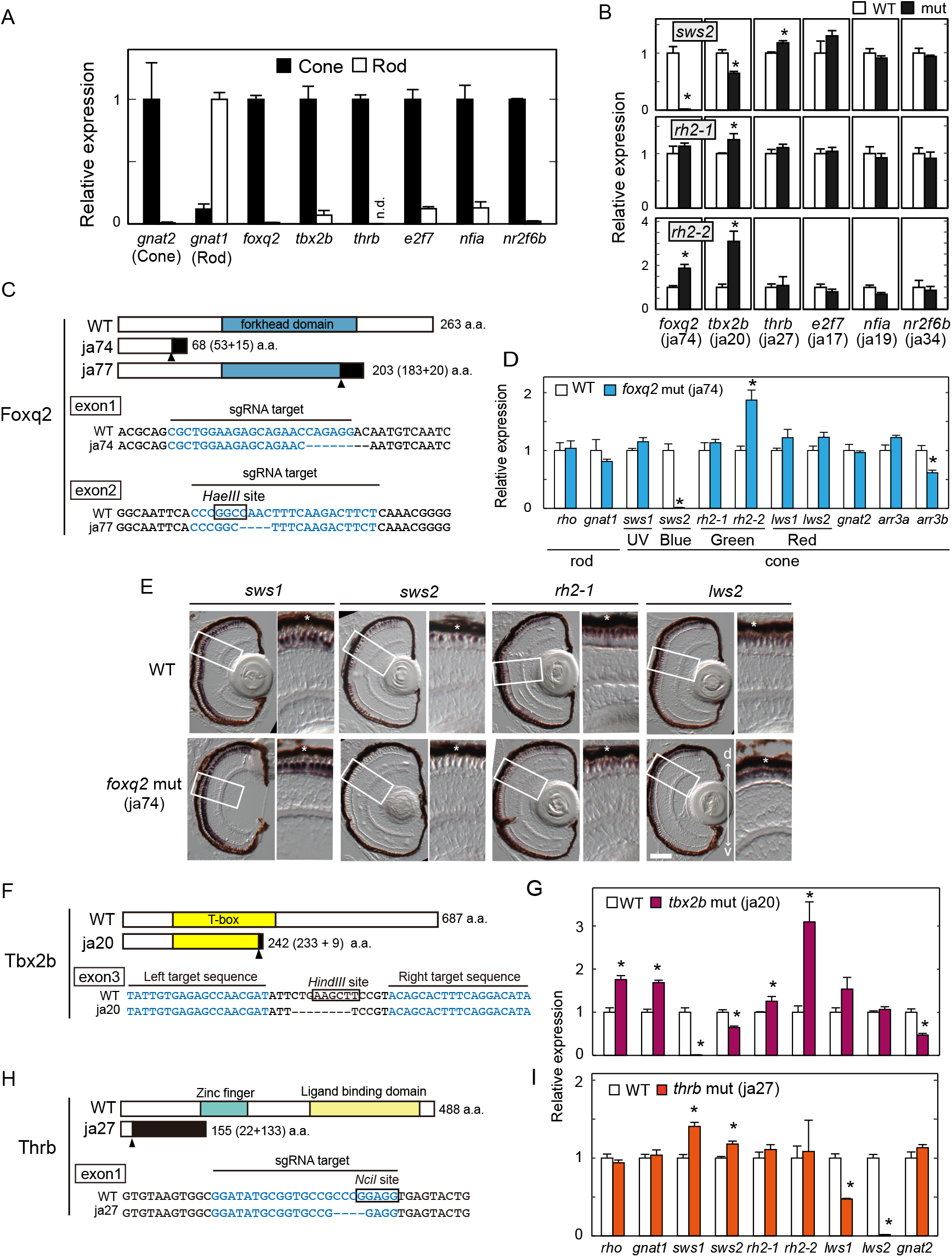
The loss of function analysis for cone-enriched transcription factors. (A) Relative expression levels of phototransduction genes and transcription factors in isolated rods and cones at the adult stage (Mean ± SD, *n* = 2). *n.d.*, not detected. (B) Relative expression levels of *sws2* opsin in the larval eyes at five days post fertilization (dpf). Mean ± SEM. \**P* < 0.05 by Student’s *t*-test. The number of fish used was as follows: *n* = 5 (*foxq2* WT), *n* = 5 (*foxq2* mut); *n* = 5 (*tbx2b* WT), *n* = 5 (*tbx2b* mut); *n* = 3 (*thrb* WT), *n* = 4 (*thrb* mut); *n* = 4 (*e2f7* WT), *n* = 4 (*e2f7* mut); *n* = 3 (*nfia* WT), *n* = 4 (*nfia* mut); *n* = 4 (*nr2f6b* WT), *n* = 4 (*nr2f6b* mut). See also Fig. S3. (C, F, H) Schematic representation of Foxq2, tbx2b and Thrb, and their partial nucleotide sequences. The frameshift site is indicated by an arrowhead. Nucleotide deletions are indicated by dashes. The nucleotide sequences (letters in blue) indicate the target sequences of TAL-effector nucleases or Cas9-sgRNA complexes. The recognition sites of the restriction endonucleases, *Nci*I, *Hind*III and *Hae*III, are surrounded by black lines. The ja74 and ja77 mutation caused a frame shift of the amino acid sequence of Foxq2 by 8-bp and 4-bp loss, respectively. The ja20 mutation caused a frame shift of the amino acid sequence of tbx2b by 8-bp loss. The ja27 mutation (4-bp loss) caused a frame shift of the amino acid sequence of Thrb2, which is an isoform of Thrb and essential for LWS opsin expression in mice and zebrafish (*15, 16*). (D, G, I) Expression profiles of phototransduction genes in the 5 dpf-larval eyes. Mean ± SEM. \**P* < 0.05 by Student’s *t*-test. The number of fish used was as follows: *n* = 5 (*foxq2* WT), *n* = 5 (*foxq2* mut); *n* = 5 (*tbx2b* WT), *n* = 5 (*tbx2b* mut); *n* = 3 (*thrb* WT), *n* = 4 (*thrb* mut). The expression levels of *sws2* and *rh2* genes are reproduced in the panel B. (E) Expression pattern of cone opsin genes examined by *in situ* hybridization using 5-dpf larval eyes of the *foxq2* mut (ja74). Magnified view (a box surrounded with white lines) is indicated in the right side of each panel. The retinal pigmented epithelium (RPE, indicated by asterisks) is adjacent to the photoreceptor layer. d-v, dorsal-ventral retina. Scale bar, 50 μm.

We generated loss-of-function mutants of zebrafish for each of the six transcription factors by introducing a frameshift mutation. In these mutants, ocular transcript levels of the middle wavelength-sensitive opsin genes were examined by RT-qPCR analysis (Fig. 1B). Among the mutant larvae, a *foxq2* mutant (Fig. 1C, ja74) displayed the most striking reduction in *sws2* expression as compared to the WT siblings (Fig. 1B). Another mutant line of *foxq2* (Fig. 1C, ja77) similarly showed a severe reduction of *sws2* expression in the larvae (Fig. S2A). The decrease of *sws2* expression in the mutant larvae (Fig. 1D and S2A) was not due to delayed development of SWS2 cones because the adult *foxq2* mutant (ja77) also exhibited minimal expression of *sws2* gene (Fig. S2B). In the *foxq2* mutant retinas, *in situ* hybridization signals of *sws2* transcripts were undetectable with no apparent change in retinal morphology at the larval (Fig. 1E) and adult stages (Fig. S2C). These results demonstrate that *foxq2* is indispensable for *sws2* expression in SWS2 cone subtype. In parallel, the *foxq2* mutation caused significant increase in mRNA levels of two major *rh2* subclasses, *rh2-1* in the larvae (Fig. S2A, ja77) and *rh2-2* in the adult (Fig. S2B, ja77), as well as upregulation of *rh2-2* in the larvae (Fig. 1D, ja74 and Fig. S2A, ja77) and *rh2-4* in the adult (Fig. S2B, ja77). The upregulation of *rh2* gene expression is likely due to transcriptional compensation for severe reduction of *sws2* expression caused by the loss of Foxq2 function.

Tbx2b is also a cone-specific transcription factor (Fig. 1A) and is required for expression of *sws1* gene (*18*). In the present study, *tbx2b ^ja20^* mutant was designed to encode C-terminally truncated Tbx2b protein (Fig. 1F), which lacked two DNA-recognition helices of T-box domain (*24*) in a manner similar to *tbx2b^fby^* mutant (*25*). As reported in *tbx2b^fby^* (*18*), *tbx2b^ja20^* larvae exhibited a severe decrease of *sws1* and a parallel increase of rhodopsin (*rho*) expression (Fig. 1F and 1G). Of note, we noticed that *tbx2b^ja20^* mutation caused 40% reduction of *sws2* expression level from that in the WT siblings and substantial increase in mRNA levels of *rh2-1* and *rh2-2* (Fig. 1G). Although the effect of *tbx2b* mutation on *sws2* expression was weaker than that of *foxq2* mutation, *tbx2b* appears to play a supportive role to *foxq2* on *sws2* expression in addition to the essential role on *sws1* expression.

Thrb is another cone-specific transcription factor (Fig. 1A), and its mutant *thrb^ja27^* (Fig. 1H) manifested a massive reduction of both *lws1* and *lws2* expression and a concomitant increase in *sws1* expression in the larval eyes (Fig. 1I) as reported in previous studies (16, *26–28*). We noticed that *thrb^ja27^* mutant exhibited a moderate but significant increase in *sws2* gene expression (Fig. 1I). The *sws2* upregulation accompanied no significant reduction of *rh2* expression level (Fig. 1I). Accordingly, Thrb contributes to fine-tuning of *sws2* expression together with a dominant role in regulation of *lws* expression. In contrast, *nfia*, *nr2f6b*, and *e2f7* mutant fish showed no noticeable change in *sws2* or *rh2* expression (Fig. 1B and S3). These three transcription factors are dispensable for middle-wavelength cone opsin gene expression.

Collectively, our mutant analysis demonstrates that *sws2* expression is predominantly regulated by a cone-specific transcription factor, Foxq2, with the aid of regulation by Tbx2b and Thrb.

### SWS2 cone subtype-specific expression of *foxq2*

To gain insights into how *foxq2, tbx2b*, and *thrb* regulate *sws2* and *rh2* expression, we investigated gene expression profiles of *foxq2, tbx2b*, and *thrb* among the four cone subtypes: SWS1, SWS2, RH2, and LWS. These four cone cells were isolated from four different lines of transgenic fish, each of which expresses EGFP in one of the four cone subtypes (Fig. 2A; see Materials and Methods for details). The cone subtype-enrichment in these purified samples was validated by RT-qPCR analyses of cone opsin genes (Fig. 2B).

**Fig. 2.**
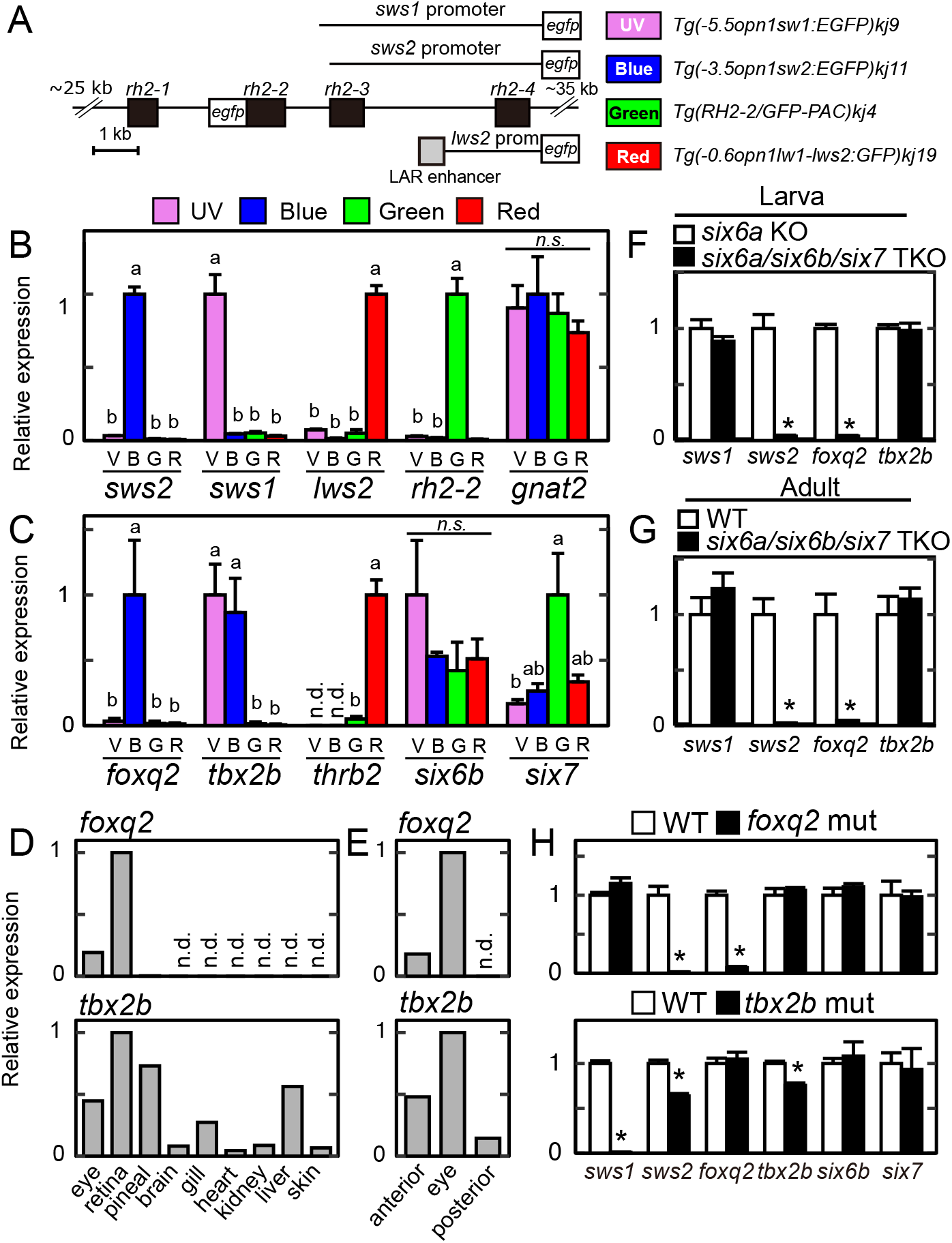
Gene expression profiles of *tbx2b* and *foxq2*. (A) Schematic drawing of transgenes for the four transgenic lines each expressing EGFP in SWS1 (UV), SWS2 (Blue), RH2 (Green) or LWS (Red) cone subtype. See also materials and methods. (B, C) Relative expression levels of cone opsin genes (B) and transcription factors (C) among isolated cone subtypes at the adult stage. Mean expression values with SEM (*n* = 3) are indicated as bars. Distinct letters indicate statistically significant differences (*P* < 0.05 by Tukey’s multiple comparison test). *n.d.*, not detected. *n.s.*, not significant. (D, E) Relative expression levels of *tbx2b* and *foxq2* in adult tissues (D), 4-dpf larval anterior segments, posterior segments and eyes (E). *n.d.*, not detected. (F) Relative expression levels of *tbx2b* and *foxq2* in the 5-dpf larval eyes of the *six6a/six6b/six7* TKO. Mean ± SEM (*n* = 5). \**P* < 0.05 by Student’s *t*-test. Note that the *six6a* KO showed similar levels of cone opsin gene expression as compared to the wild-type [See Ref. (*23*)]. (G) Relative expression levels of *tbx2b* and *foxq2* in the adult eyes of the *six6a/six6b/six7* TKO. All of the fish used here are in the transgenic background, *Tg(−3.5opn1sw2:EGFP)^kj11Tg^*, where EGFP is expressed in the SWS2 cone subtype. Data are represented by mean ± SD (*n* = 3, WT; *n* = 2, *six6a/six6b/six7* TKO). (H) Relative expression levels of cone opsins and transcription factors responsible for photoreceptor gene expression in the eyes of the *tbx2b* (ja20) and *foxq2* (ja74) mutants. Mean ± SEM. \**P* < 0.05 by Student’s *t*-test. The number of fish used was as follows: *n* = 5 (*tbx2b* WT), *n* = 4 (*tbx2b* mut); *n* = 5 (*foxq2* WT), *n* = 5 (*foxq2* mut).

Subsequent analysis of *foxq2, tbx2b*, *thrb, six6b*, and *six7* in the four samples revealed that *foxq2* was specifically expressed in SWS2 cone subtype (Fig. 2C). Another SWS2 regulator, *tbx2b*, was expressed in both SWS1 and SWS2 cone subtypes, whereas *thrb* was expressed only in the LWS cone subtype (Fig. 2C). In contrast, SWS2/RH2 regulators, *six6b* and *six7*, were expressed in all the cone subtypes (Fig. 2C). The SWS2 cone subtype-enriched expression of *foxq2* and *tbx2b* suggests that these two factors coordinately regulate the cell-type-specific expression of *sws2.*

*foxq2* expression was restricted to the retina among various adult tissues (Fig. 2D), and in the larvae, it was detected only in the anterior half of the body (including the eyes) but not in the posterior half (Fig. 2E). These observations suggest that in the course of development, *foxq2* expression becomes restricted to SWS2 cones within the retina. Importantly, the mRNA level of *foxq2* was severely reduced in *six6a/six6b/six7* triple knockout (TKO) lacking *sws2* expression, *i.e.*, loss of SWS2 cone identity from the retina (*23*) (Fig. 2F and G). *foxq2* expression was also abrogated in the larval eyes of *foxq2* mutant (ja74) as compared to that of WT sibling (Fig. 2H). The close correlation between *foxq2* and *sws2* expressions imply that Foxq2 is responsible for establishing SWS2 cone identity. Meanwhile, *tbx2b* was expressed widely in various tissues at the larval (Fig. 2E) and adult stage (Fig. 2D), and the ocular *tbx2b* expression level was unaffected in the *foxq2* mutant (Fig. 2H) and *six6a/six6b/six7* TKO (Fig. 2F and 2G), both of which were deficient in *sws2* expression. These observations indicate dominant expression of *tbx2b* outside of SWS2 cones and imply that Tbx2b may have pleiotropic roles for eye development (*29*) including the cone identity determination.

### Functional interaction of Foxq2 with *sws2* promoter

We then investigated functional interaction of Foxq2 with *sws2* promoter. Foxq2 protein (Fig. 1C) has a conserved DNA-binding domain, termed forkhead domain composed of about 100 amino acid residues (*30*). In Fox transcription factor family, Foxq2 is categorized into clade I forkhead proteins (*31*), which recognize two types of common forkhead-target DNA motifs termed FkhP (RYAAAYA) and FkhS (AHAACA) (*32*). Our motif scanning analysis (see Materials and Methods) revealed two FkhP motifs and two FkhS motifs present within 1.56-kb *sws2* promoter region (Fig. 3A and S4), which drives selective gene expression in SWS2 cones (*33*). In a cell-based reporter assay, the 1.56-kb *sws2* promoter was transactivated by VP64-Foxq2 (Fig. 3B and 3C), in which Foxq2 is N-terminally fused with four repeats of the VP16 transcriptional activator domain [Fig. 3B and (*34*)]. The transcriptional activation was attenuated by deletion of the upstream region of 1.03 or 1.26 kbp (0.53-kb or 0.3-kb *sws2* promoter, respectively), leaving single FkhS motif (Fig. 3A and 3C). Still, we observed more than 30-fold transactivation of the 0.3-kb *sws2* promoter by VP64-Foxq2, while the activation was largely reduced by the complete deletion of the promoter sequence (Fig. 3C). The VP64-Foxq2-dependent transactivation of a shorter promoter (0.25-kb *sws2* promoter) was markedly decreased by introducing a 6-bp mutation in the FkhS motif (Fig. 3D). We then generated a mutant protein, termed Fork-del, lacking nine amino acid residues that are required for specific DNA-binding of the forkhead domain (*32*) (Fig. 3B). The protein mutation abolished the ability of the transcriptional activation of the *sws2* promoter (Fig. 3B, 3D and 3E). These results demonstrated that Foxq2 functionally interacted with the forkhead-target DNA motif in the *sws2* promoter. Together with the SWS2-specific expression of *foxq2* (Fig. 2), Foxq2-mediated transcriptional regulation of *sws2* gene would reasonably account for the selective expression of *sws2* opsin in SWS2 cone subtype.

**Fig. 3.**
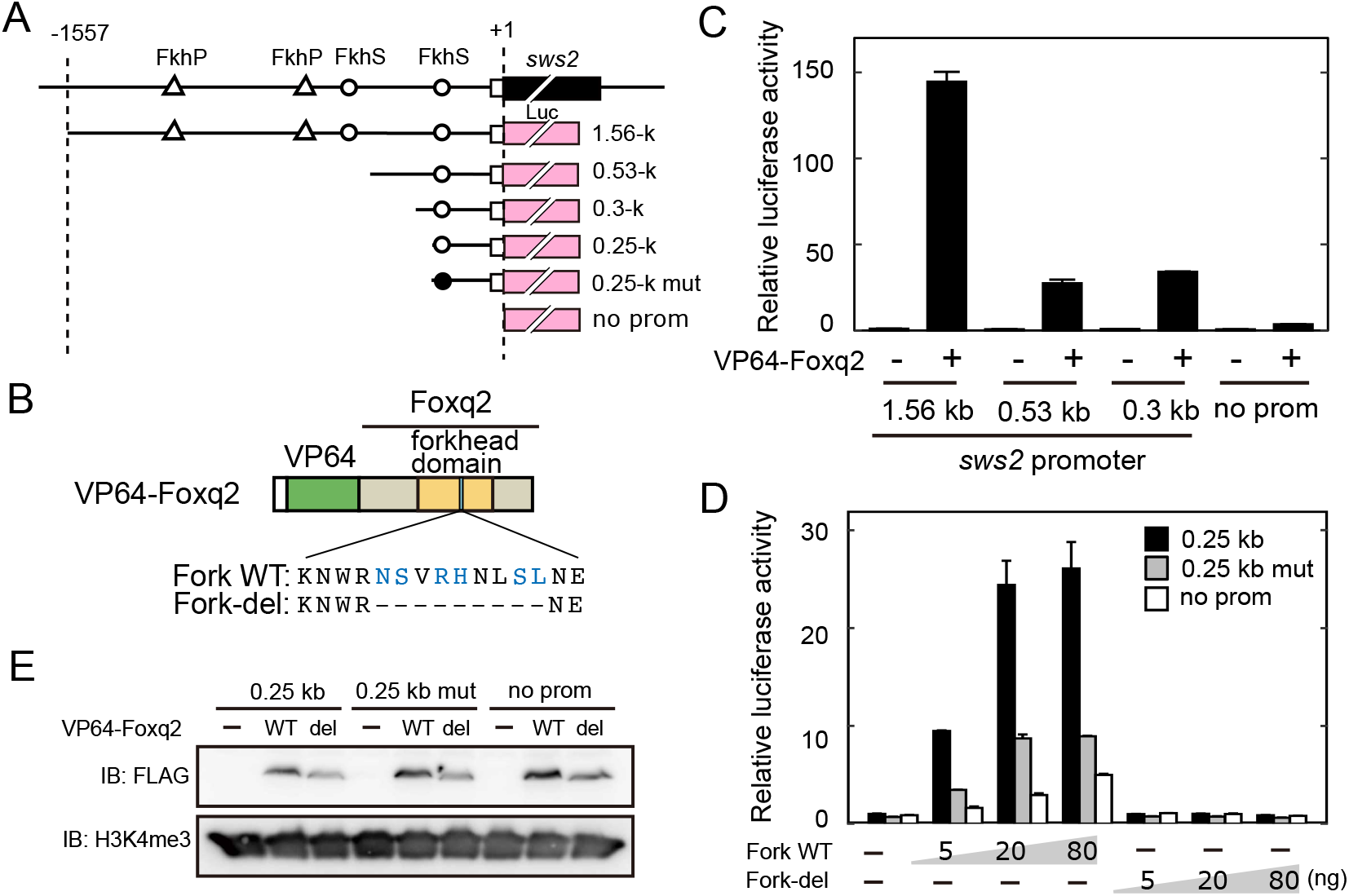
DNA binding of Foxq2 to *sws2* promoter. (A) Schematic structure of the promoter-luciferase reporter constructs for zebrafish *sws2*. The coding (solid box) and the untranslated (open box) regions of the first exon are indicated. The translational start site is marked as +1. The forkhead binding motif (ACAACA) and its mutant (GTGGTG) are marked as open and closed circles, respectively. See also Fig. S4. (B) Schematic representation of VP64-Foxq2 protein. The Foxq2 mutant protein (Fork-del) lacks the canonical Fox base-contacting residues in the forkhead domain. These critical amino acid residues are highlighted in blue according to the previous paper (*32*). (C, D) The transcriptional assay in HEK293T17 cells using the luciferase reporter containing the promoter region of zebrafish *sws2*. The luciferase activity derived from the firefly luciferase reporter was normalized to that from the Renilla luciferase reporter. Data are represented by the mean ± SD (*n* = 2). (C) The luciferase activity in each condition was normalized to the mean value obtained from the cells transfected with both empty vector and 1.56-kb *sws2* reporter vector (leftmost bar). VP64-Foxq2 expression plasmid (100 ng) was used for the transfection. (D) The amount of VP64-Foxq2 expression plasmid used for each well is indicated in the graph. The luciferase activity in each condition was normalized to the mean value obtained from the cells transfected with both the empty expression vector and the 0.25-kb *sws2* (0.25 kb) reporter vector (leftmost bar). (E) Protein expression of VP64-Foxq2 (WT) and its mutant protein (Fork del) in the HEK293T17 cells transfected with 80 ng of the expressing plasmid. The antibody against a histone modification (H3K4me3) is served as a loading control.

### Fate determination of SWS2 cone subtype governed by *foxq2*

The *foxq2* mutants (ja74 and ja77) were not only deficient in *sws2* expression but also characterized by significant reduction in mRNA level of arrestin 3b (*arr3b*), a cone phototransduction gene selectively expressed in SWS1 and SWS2 cone subtypes (*35*) (Fig. 1D and S2). These observations suggest that *foxq2* deficiency impairs development of SWS2 cone cells and/or their maintenance. To gain further insight into the fate determination of SWS2 cone subtype, we generated *foxq2* transgenic (*foxq2-tg*) zebrafish, *Tg(−5.2crx:EGFP-2A-FLAG-foxq2)*, in which expressions of both EGFP and Foxq2 are driven by 5.2-kb *crx* promoter in all the developing and matured photoreceptor cells (Fig. 4A and 4B) (*16*). RT-qPCR analysis of larval ocular mRNAs in three *foxq2-tg* lines (ja78Tg, ja79Tg, and ja91Tg) revealed that forced expression of *foxq2* in the wild-type background caused a severe decrease in *sws1* mRNA level (Fig. 4C and S5). This phenotype was not accompanied by any detectable change in the expression levels of *tbx2b* and *crx* (Fig. 4D and S5), which are regulators of *sws1* expression (*18*) (Fig. 1G). It is most probable that Foxq2 suppresses *sws1* expression in cones (Fig. 4E).

**Fig. 4.**
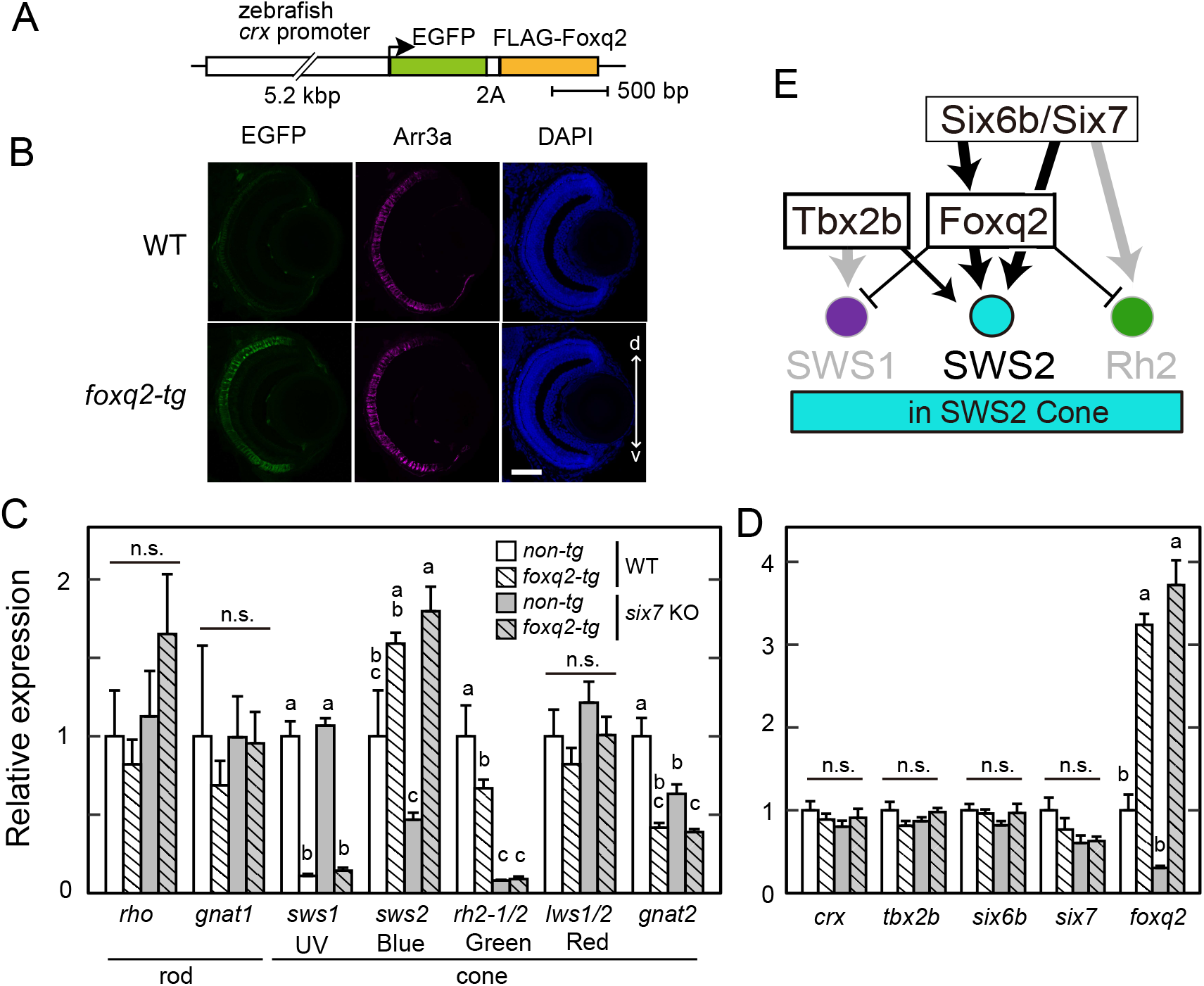
Foxq2-mediated transcriptional regulation of *sws2* downstream of Six7. (A) Schematic drawing of a transgene construct used for generating *foxq2*-tg. (B) The immunofluorescent image in the *foxq2-tg* (ja78Tg) larvae at 5-dpf. Scale bar, 50 μm. d-v, dorsal-ventral retina. (C, D) Relative mRNA levels of opsin genes and transcription factors in the 5-dpf larval eyes of the *foxq2-tg* (ja78Tg) and/or *six7* KO (mean ± SEM). Distinct letters indicate statistically significant differences (*P* < 0.05 by Tukey’s multiple comparison test). *n.s.*, not significant. The number of fish used was as follows: *n* = 3 (WT), *n* = 5 (*six7* KO), *n* = 4 (*foxq2-tg*) and *n* = 5 (*foxq2-tg/six7* KO). (E) Hypothetical model of transcriptional network accounting for *sws2* opsin expression in the SWS2 cone subtype. Transcriptional regulation in the SWS2 cone subtype is highlighted in black, while transcriptional regulation in the other cone subtypes is highlighted in grey.

The selective expression of *foxq2* in SWS2 cones (Fig. 2C), together with binding of Six6b and Six7 to *foxq2* gene locus (Ref. 23 and Fig. S1), suggest that *sws2* expression is regulated by Foxq2 downstream of Six6b and Six7 (Fig. 4E). This possibility was explored by overexpressing *foxq2* in the homozygous *six7* knockout (KO) (*19*), in which expression levels of *sws2* and *foxq2* are markedly reduced (Fig. 4C and 4D). The forced expression of *foxq2* in the *six7* KO background (*foxq2-tg;six7* KO) recovered the reduced expression of *sws2* up to a level higher than that in the WT control (Fig. 4C). In contrast, the overexpression of *foxq2* had no significant effect on the severely reduced expression levels of *rh2* (*rh2-1* plus *rh2-2*) (Fig. 4C) or on mRNA levels of *crx, tbx2b, six6b*, and *six7*, transcriptional regulators of cone opsin expression (Fig. 4D). These results demonstrated that Foxq2 regulates *sws2* expression downstream of Six7, suggesting that Foxq2 is a terminal selector for SWS2 subtype identity (Fig. 4E).

### Conservation of *FOXQ2* gene among vertebrate species

*FOXQ2* gene is annotated in the genomes of a wide range of vertebrate species (Fig. 5), such as ray-finned fish (spotted gar, zebrafish, and medaka), a lobe-finned fish (coelacanth), and an avian (sparrowhawk), all of which retain *SWS2* gene. We found highly conserved gene synteny around *FOXQ2* locus among the vertebrates, and the synteny analysis of the human genome indicated the absence of any gene homologous to *FOXQ2* between *PIAS4* and *ZBTB7A* gene loci (Fig. 5B). Intriguingly, BLAST searching and subsequent manual annotation revealed that a mammalian species, platypus, retains a gene orthologous to *FOXQ2* harboring the forkhead domain highly conserved among FOXQ2 subfamily members (Fig. 5 and S6; see also supplementary text). The oviparous mammals including platypus diverged from marsupial and placental mammals at the earlier stage of mammalian evolution (around 200 million years ago). Of note, platypus retains *SWS2* gene in its genome (*36*), whereas many other mammals have lost it (*7, 37*) most likely due to a long evolutionary history of nocturnality in mammalian ancestors (*38*). These lines of genomic evidence suggest that Foxq2-dependent *SWS2* expression is a highly conserved regulatory mechanism that was acquired at the early stage of vertebrate evolution.

**Fig. 5.**
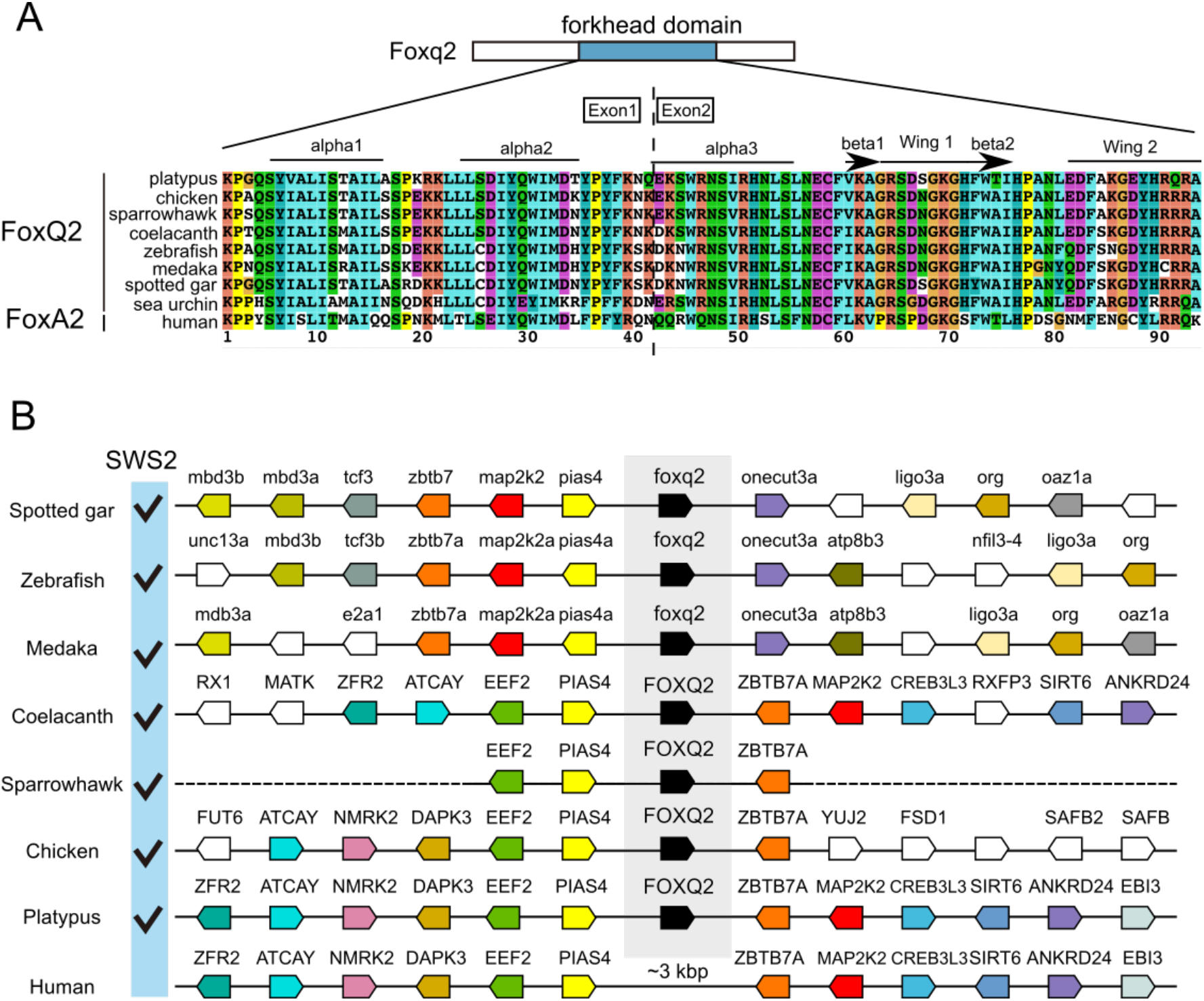
*FOXQ2* gene conservation among animal species. (A) Sequence alignment of forkhead domain of FOXQ2 and FOXA2 proteins. Secondary structure elements are indicated at the top according to the FOXA2-DBD/DBE2 complex in the previous study (*69*). (B) Genomic environment of *FOXQ2* genes in vertebrate species. Orthologous genes contributing to the conserved order of *FOXQ2* genes among vertebrates are similarly color-coded. The presence of *SWS2* gene in each species is indicated as checkmarks according to the previous study (*70*). Accession numbers for the genomic location of *FOXQ2* in each species is indicated in Table S5.

## Discussion

Under the solar light irradiation, both terrestrial and aquatic environments on the earth’s surface are enriched with blue-to-green region of visible spectrum (*39*), which is detected by cone photoreceptor cells expressing middle-wavelength sensitive opsin genes, SWS2 and RH2. Their expression requires transcription factors, Six6 and Six7, in zebrafish (*19, 23*), and the *six6/six7* mutant fish lacking both SWS2 and RH2 cones show severely reduced survival rate due to impairment of visually driven foraging behavior (*23*). A recent study of *in vivo* calcium imaging of zebrafish cone photoreceptors reported that SWS2 and RH2 cones, but not SWS1 or LWS cones, display strong spectral opponency and efficiently extract chromatic information from the natural light spectrum (*40*). In this way, a combination of spectrally distinctive SWS2 and RH2 cones particularly plays an important role in the tetrachromatic visual system. The present study explored the transcriptional regulatory logic that defines SWS2 and RH2 cone identities. For this purpose, cone-enriched transcription factors were comprehensively identified by the transcriptome analysis with purified cone cells (vs. purified rod cells; Data S1). Subsequent functional analyses of these transcription factors demonstrated that Foxq2 is indispensable for *sws2* expression (Fig. 1). We pursued expression profiling of the cone-enriched transcription factors among the four isolated cone subtypes (Fig. 2A–C). *foxq2* is selectively expressed in SWS2 cone, whereas all cone subtypes express *six6b* and *six7* (Fig. 2C), which are required for both *sws2* and *rh2* gene expression (*19*). A transcriptional network is deciphered in which Foxq2 acts as a downstream regulator of Six7 (Fig. 4A–D) and regulates *sws2* expression (Fig. 3). A wide range of vertebrate species retain both *FOXQ2* and *SWS2* gene (Fig. 5). These lines of evidence demonstrate that Foxq2 determines SWS2 cone identity in zebrafish (Fig. 4E) and suggest that Foxq2-dependent *sws2* expression is a highly conserved regulatory mechanism that was acquired at the early stage of vertebrate evolution.

Quantitative comparison of transcription factor expression among the four isolated cone subtypes (Fig. 2C) provides valuable information about terminal differentiation of each cone subtype in zebrafish. *foxq2*, a crucial regulator of *sws2* expression, is selectively expressed in SWS2 cones, indicating that Foxq2 is a *bona fide* terminal selector of SWS2 cone. On the other hand, *tbx2b*, known as a master regulator of SWS1 cone (*18*), is expressed in not only SWS1 cones, but also SWS2 cones (Fig. 2C), in which Tbx2b acts as a supportive regulator of *sws2* expression (Fig. 1G). The expression of the SWS1 master regulator in SWS2 cones implies that some mechanism should suppress *sws1* misexpression in SWS2 cones. It is probable that Foxq2 contributes to the *sws1* suppression because the forced expression of *foxq2* markedly reduced *sws1* expression in SWS1 cones (Fig. 4C and S5). Similarly, Foxq2 appears to suppress *rh2* misexpression in SWS2 cones (Fig. 1D, 4C, and S2). Thus, Foxq2 has dual functions acting as an activator of *sws2* transcription and as a suppressor of *sws1* and *rh2* genes in SWS2 cones (Fig. 4E) and, presumably, in developing cone cells. The dual functions of Foxq2 would enhance robustness of SWS2 cone identity. Meanwhile, *thrb* is predominantly expressed in LWS cone subtype (Fig. 2C), being consistent with the widely accepted idea that Thrb is a master regulator for the LWS cone (*15–17*). Collectively, Foxq2, Tbx2b, and Thrb can be defined as the terminal selectors (*41*) that govern cell fate determination of SWS2, SWS1, and LWS cone subtypes, respectively.

In contrast to these three terminal transcriptional selectors, *six7* is widely expressed among the four cone subtypes (Fig. 2C) and thus *six7* is unlikely to be a terminal selector of RH2. Although Six7 is indispensable for expression of *rh2* (*rh2-1, rh2-2, rh2-3*, and *rh2-4*; Ref. 19 and Fig. 4C), terminal differentiation of RH2 cone might be mediated by an unidentified transcription factor. An alternative idea is that Rh2 cone identity is established by both (i) the presence of *six7* and (ii) the absence of the terminal selectors such as *tbx2b* (for SWS1), *foxq2* (for SWS2), and *thrb* (for LWS). In this scenario, differentiation toward Rh2 cones should be a default pathway in the cone development, because Rh2 cone differentiation is governed by *six7* (*19*) and *six7* expression begins as early as that of *crx*, a master regulator of cone and rod development (19, 39). It is worth noting that the “Rh2-default” hypothesis is consistent with molecular phylogeny of cone opsin genes, in which RH2 subfamily diverged from SWS2 at the latest step of the molecular evolution of the cone opsins (4, 5, 40). Building on these assumptions, we propose that the tetrachromatic color vision system in ancestral vertebrates employed the “RH2-default” mechanism of cone differentiation as is observed in zebrafish. In mammals, on the other hand, RH2 and SWS2 genes have been lost, and hence it is likely that the “RH2-default” mechanism has been modified to a “SWS1-default” mechanism. In the latter mechanism, the presence or absence of only one terminal selector, THRB, directs differentiation between LWS and SWS1 cones as shown in mice (*8, 15*).

*FOXQ2* gene is evolutionarily conserved not only among vertebrates but also in many invertebrates (*31, 32, 44*). In embryos of the invertebrates, *FOXQ2* is responsible for specification and positioning of anterior neuroectoderm (*45, 46*). The anterior neuroectoderm develops into central nervous system, in which a subset of sensory cells express a photoreceptive protein, c-opsin, that is orthologous to vertebrate cone and rod opsins (*47, 48*). FOXQ2 might regulate differentiation of the c-opsin-expressing cells from progenitor cells in the anterior neuroectoderm. Intriguingly, a transcriptional network in the anterior neuroectoderm of invertebrates also includes *six3*, a gene orthologous to vertebrate *six3*, *six6* and *six7* (*45, 46*). The presence of the Six3/6/7-Foxq2 transcriptional network in both vertebrate and invertebrate species suggest that their common ancestor employed this transcriptional network for development of the light-sensitive cells. In the vertebrate lineage, the Six3/6/7-Foxq2 transcriptional network might have been co-opted for establishment of SWS2 cone identity during or after the appearance of a full set of cone opsins, thereby conferring high-acuity discrimination of blue-to-green region of visible light.

## Materials and Methods

### Zebrafish

The Ekkwill strain was used as the wild-type zebrafish. All appropriate ethical approval and licenses were obtained from Institutional Animal Care and Use Committees of The University of Tokyo. All the procedures were conducted according to local guidelines of the University of Tokyo. Zebrafish were raised in a 14-h light/10-h dark cycle and fed twice per day with live baby brine shrimps. Embryos were raised at 28.5 °C in egg water (artificial seawater diluted 1.5:1,000 in water). Mutant strains and transgenic lines used in the present study are listed in Table S1.

### Purification of rod and cone photoreceptor cells

For isolation of rod and cone photoreceptor cells, we used the transgenic zebrafish lines, *Tg(rho:egfp)^ja2^* (*49*) and *Tg(gnat2:egfp)^ja23^* (*19*), which express EGFP in rods and all cone subtypes, respectively. The rod and cone cells were isolated as described previously (*19, 23, 50*). Briefly, retinas were dissected from dark-adapted adult fish under dim red light. The isolated retinas were digested with 0.25% trypsin, 10 U/ml DNaseI, 2 mM MgCl_2_, and 2 mM EGTA in Ca^2+^-free Ringer’s solution for 30 min at 37°C. The reaction was terminated by adding soybean trypsin inhibitor (final 0.5%) and fetal bovine serum (final 10%). The dissociated cells were filtrated through a 35-μm nylon mesh (Falcon). EGFP-positive cells were isolated with a fluorescence activating cell sorter (FACSAria, BD Biosciences) by the following three parameters: forward scatter, side scatter and green fluorescence. The isolated cells were directly collected into 800 μL of TRIzol reagent (Thermo Fisher Scientific) in 1.5 ml microtubes.

For isolation of cone subtypes, we used four lines of transgenic zebrafish: *Tg(−5.5opn1sw1:EGFP)^kj9^* (*51*), *Tg(−3.5opn1sw2:EGFP)^kj11^* (*33*), *Tg(RH2-2/GFP-PAC)^kj4^* (*52*) and *Tg(−0.6opn1lw1-lws2:GFP)^kj19^* (*53*), which express EGFP in SWS1, SWS2, RH2 and LWS cone subtypes, respectively. The cone photoreceptor isolation was carried out as described above with some modifications as follows. The isolated cells were directly collected into 450 μL of the lysis buffer (RNeasy Lysis Buffer, Qiagen) in 1.5 ml microtubes.

### Microarray

Total RNA was isolated from the sorted cells using RNeasy Extraction kit (Qiagen). Quality and quantity of the resulting RNA were assessed using a NanoDrop ND-2000 spectrophotometer (Thermo Scientific) and an Agilent 2100 Bioanalyzer (Agilent Technologies). Microarray Analysis was performed using the Agilent 4 ×44 k Zebrafish microarray according to the manufacturer’s protocol for the two-color method. Cy3- or Cy5- labeled cRNA probe was synthesized from 150 ng of total RNA using the Quick-Amp Labeling Kit (Agilent Technologies). Quantity of the resulting labeled cRNA was assessed using a NanoDrop ND-2000 spectrophotometer. Equal amounts of Cy3 and Cy5-labeled cRNA (825 ng) from two different samples were hybridized to zebrafish microarrays (Agilent Zebrafish Oligo Microarrays ver.2, G2519F-019161) for 17 h at 60 °C. In order to compare gene expression profiles between cone and rod samples, we analyzed two independent biological replicates as following: (1) Cy3-rod#1 cRNA and Cy5-cone#1 cRNA and (2) Cy5-rod#2 cRNA and Cy3-cone#2 cRNA. The hybridized arrays were then washed and scanned using an Agilent microarray scanner (G2505 C; Agilent Technologies). Data were extracted from the scanned image using Feature Extraction version 10.5.1.1 (Agilent Technologies). We then listed differentially expressed genes with Microsoft Excel as follows: 1) We excluded any probes whose signals in two arrays were all determined as negative, meaning that probes were not flagged as “WellAboveBG” in any of two arrays. 2) We selected all the probes whose signal intensities vary largely between the photoreceptor cell types according to the following thresholds of the averaged ratios: 10-fold increase for cone compared with rod and 4-fold increase for rod compared with cone. 3) We checked whether both of the two independent biological replicates showed similar expression profiles, and selected probes whose absolute values of the ratios in the two biological replicates were both >2.0 for rod/cone or both >3.0 for cone/rod. The lower threshold was given for the rod/cone ratio because a small but noticeable level of rod contamination was detected in cone samples such as *gnat1* (Fig.1A). We set these threshold values based on the ratios observed for known cone-specific genes. The microarray datasets will be available at the National Center for Biotechnology Information Gene Expression Omnibus (GEO) database.

### Generation of mutant zebrafish

The *foxq2*, *thrb*, and *nr2f6b* mutants were generated with a CRISPR/Cas9 system. The sgRNA sequences (Table S2) were designed to target exon 1 or exon 2 (Fig. 1C, 1H, and S3E) using CRISPRdirect (https://crispr.dbcls.jp/) (*54*), CHOPCHOP (https://chopchop.cbu.uib.no) (*55*), or CRISPRscan (*56*). The sgRNA was synthesized by a cloning-free method as previously described (*57*). For generating Cas9 mRNA, the template plasmid DNA, pCS2-nCas9n (addgene #47929) (*58*) or pCS2+hSpCas9 (addgene #51815) (*59*) were used for *in vitro* transcription. Cas9 mRNA was synthesized with the SP6 mMESSAGE mMACHINE Kit (Ambion) and purified with the RNeasy Mini kit (Qiagen). The solution containing 200-pg Cas9 mRNA and 25-pg sgRNA was injected into the cytoplasm of the one cell-stage embryos.

The *tbx2b*, *e2f7*, and *nfia* mutants were generated by TALENs as previously described (*19, 23*). To target each of the three genes, a pair of the TAL effector repeats (Table S2) were designed to target exon 2 or exon 3 with the Golden Gate assembly methods (Fig. 1, S3A, and S3C). TALEN mRNA was synthesized using the SP6 mMESSAGE mMACHINE Kit (Ambion) and purified with the RNeasy Mini kit (Qiagen). The solution containing 200-pg each of the two TALEN mRNAs was injected into the cytoplasm of the one cell-stage embryos.

The injected fish (F0) were crossed with the wild-type zebrafish. The resultant F1 offspring were screened for the presence of CRISPR/Cas9- or TALENs-induced mutations by a combination of PCR and subsequent enzyme digestion for *thrb, nr2f6b, tbx2b, e2f7*, and *nfia* mutants, or by Heteroduplex Mobility Assay (HMA) for *foxq2* mutants as described previously (*60*). To sequence-verify mutations, genomic sequences surrounding the mutations were amplified by nested PCR, and the resultant PCR products were sequenced directly. After isolation of mutant zebrafish and verification of the mutations, the mutant genotypes were confirmed by a combination of PCR and subsequent enzyme digestion except for the *foxq2* (ja74) gene locus. The *foxq2^ja74^* mutant genotype was determined by PCR with two pairs of primers: (i) foxq2_Fw1A (5’- TGGCT AAACG AACAA ACACG -3’) and foxq2_Rv1WT (5’- ATGGA TTGAC ATTGT CCTCT G -3’); (ii) foxq2_Fw1mutA (5’- GCTGG AAGAG CAGAA CAATG -3’) and foxq2_Rv1A (5’- GGAAA TGAGG GCAAT GTAGG -3’). PCR primers used for amplification are listed in Table S3.

The mutant larvae and its siblings were dissected into anterior and posterior segments; the posterior parts were used for genotyping, while the anterior segments were soaked in RNA*later* (Sigma) for RT-qPCR analysis or fixed with 4% paraformaldehyde (PFA) in Ca^2+^- and Mg^2+^-free Dulbecco’s PBS (D-PBS) for cryosectioning.

### RT-qPCR analysis

Zebrafish were anesthetized by chilling on ice, and their tissues were collected during the light phase of the light-dark cycle and soaked in RNA*later*. After genotyping described in the previous section, the larval eyes were isolated with a needle. A pair of two larval eyes or one adult eye was considered a biological replicate for each genotype. RNA extraction and reverse transcription were conducted as described previously (*19, 23*). In short, RNA was extracted and purified with RNeasy Micro Kit or RNeasy Mini Kit (Qiagen). In all the experiments except for Fig. 1A, the extracted RNA was reverse-transcribed into cDNA with the oligo (dT)_15_ primer with GoScript^TM^ Reverse Transcriptase (Promega). In Fig. 1A, the reverse transcription was conducted with SuperScript II (Thermo Fisher Scientific) using anchored (dT)_16_ primers. The reverse-transcribed cDNA was subjected to quantitative PCR using GoTaq qPCR Master Mix (Promega) and the StepOnePlus™ Real-time PCR system (Applied Biosystems) following the manufacturers’ protocols. Expression levels were calculated by the relative standard curve method. The standard curve was prepared with serial dilutions of cDNA samples reverse-transcribed from total RNA of zebrafish eye. The transcript levels were normalized to beta-actin 2 (*actb2*) transcript levels in all the figures. Primers used for quantitative PCR are listed in Table S4 and in our previous studies (*19, 23*). Total transcript level of RH2 (*rh2-1* and *rh2-2*) or LWS (*lws1* and *lws2*) opsin genes at the larval stage was measured with a set of PCR primer, which amplifies both *rh2-1* and *rh2-2* opsin genes (referred to here as *rh2-1/2*) or both *lws1* and *lws2* opsin genes (referred to here as *lws1/2*). Expression levels of all the transcript isoforms of *thrb* were measured in Fig. 1A, while in the rest of the experiments, we measured expression levels of a transcript isoform of *thrb, thrb2*, which is essential for *lws* expression in mouse and zebrafish (*15, 16*).

### *In situ* hybridization

*in situ* hybridization using ocular sections was carried out as described previously (*19, 23*). In short, zebrafish were anesthetized by chilling on ice and subjected to dissection the light phase of the light-dark cycle. The larval anterior segments or the adult eyes were fixed in 4% PFA in D-PBS overnight at 4 °C. Before the fixation, the adult eyes were enucleated and poked with tweezers to make a tiny hole in the cornea. After sucrose infiltration and OCT compound embedding, the 10-μm frozen ocular sections were prepared with a cryostat. The cryosections were pre-treated with proteinase K and hybridized with DIG-labelled cRNA probes, and the hybridization signals were visualized by NBT/BCIP staining. The images were acquired with an upright microscope (Axioplan2, Carl Zeiss). The cRNA probes were generated as described in our previous study (*19, 23*).

### Immunohistochemistry

Immunohistochemistry with ocular sections was carried out as described previously (*19, 23*). Briefly, ocular cryosections were prepared as described in the previous section. The cryosections were pre-treated with a blocking solution and then incubated with a primary antibody diluted in the blocking solution overnight at 4°C. After washed with PBS (10 mM Na-phosphate buffer, 140 mM NaCl, 1 mM MgCl_2_, pH 7.4), the treated sections were immersed again with the blocking solution, and then incubated for 4 hr at room temperature with a secondary antibody and with DAPI (3 μg/ml) for staining of the cell nuclei. The stained sections were coverslipped with VECTASHIELD Mounting Medium (Vector Laboratories) and imaged with a confocal laser scanning microscope (TCS SP8, Leica). The primary antibody used is mouse monoclonal antibody Zpr1 (diluted 1:400, Zebrafish International Resource Center, Eugene) against arrestin 3a. The secondary antibody used is goat anti-mouse IgG antibody conjugated with Alexa-568 (diluted to 2 μg/ml, A-11004, Molecular Probes).

### Luciferase assay

For constructing the firefly luciferase reporter plasmids, the 1.56-kb *sws2* upstream region with its 5’-UTR was amplified by PCR. The amplified fragment was ligated using the In-Fusion cloning kit (Takara) into the pGL4.13[luc2/SV40] (E6681, Promega) digested with *Hind*III and *Bgl*II. PCR primers used were as follows: IF-sws2-1.5kFwBgl (5’- CGAGG ATATC AGATC TAACG ATGTT TGCTG TTTGT TC -3’) and IF-sws2-RvHind (5’- CCGGA TTGCC AAGCT TCTTG CTTGT AATTG GTGCC C -3’). The 0.53-kb *sws2* reporter was constructed in a manner similar to the 1.56-kb *sws2* reporter with following primers: IF_532_sw2_FwXho (5’- GCTCG CTAGC CTCGA GCAAC TCTCA AGTAT TTAAG G -3’) and IF-sws2-RvHind. The 1.56-kp *sws2* reporter was truncated by PCR to generate the 0.3-kb and 0.25-kb *sws2* reporter by using the following PCR primers: sws2_300_Fw (5’- TCTTG TACTG CGCAG ATGTA G -3’), 250sw2_Fw (5’- GAAAC TTTGT GTGTA GCTGA TG -3’), and pGL4.13_Rv (5’- AGATC TGATA TCCTC GAGGC TAG -3’). For generating the pGL4 vector having no basal promoter, the SV40 promoter in the pGL4.13 vector was removed by a combination of enzyme digestion and self-ligation; the pGL4.13 vector was double digested with *Hind*III and *Xho*I, treated T4 DNA polymerase to make a blunt end, and self-ligated. The nucleotide mutations on the potential Foxq2-binding motif were introduced into the 0.25-kb SWS2 reporter by PCR. The promoter sequence in each of these resultant constructs was sequenced to confirm that no unintended mutation was incorporated into the promoter region.

To generate the expression plasmid of Foxq2, we first inserted a FLAG epitope tag into the pCAG vector (gifted from Dr. Takahiko Matsuda). The resultant vector was named as pCAG-FLAG. We then amplified the cDNA fragments of *foxq2* from retinal cDNAs of adult zebrafish, and cloned them into the *EcoR*V-treated pCAG-FLAG vector. These plasmids was named as drFoxq2/pCAG-FLAG. For generating the VP64-Foxq2 expression plasmid, we first cloned the multiple repeats of the herpes simplex VP16 activation domain (synthesized DNA fragments) into the *Xho*I-treated pCAG-FLAG vector. The resultant plasmid, named as pCAG-FLAG-VP64N, was treated with the *EcoR*V and ligated with the PCR-amplified cDNA fragment. The nucleotide deletion in the DNA binding domain of Foxq2 was introduced by PCR. The DNA sequence data for drFoxq2/pCAG-FLAG-VP64N will be available in the DDBJ/EMBL/NCBI.

HEK293T17 cells were grown in the Dulbecco’s modified Eagle’s medium supplemented with 10% fetal bovine serum, 100 units/mL penicillin, and 100 μg/mL streptomycin. HEK293T17 cells in 24-well plates were transiently transfected with polyethyleneimine (Polysciences, #24765). The firefly luciferase plasmid harboring the *sws2* promoter described above was used as a reporter plasmid (10 ng per well), while the Renilla luciferase plasmid, pGL4.74[hRluc/TK] (E6921, Promega), was simultaneously transfected as an internal control reporter (0.5 ng per well). The amount of the expression plasmid of Foxq2 used for the transfection is indicated in the figure legends. The total amount of DNA transfected in each well was equally adjusted by adding the empty expression plasmids. The transfected cells were collected 36-48 h after the transfection, rinsed with PBS(-) (137 mM NaCl, 2.69 mM KCl, 5.5 mM Na-_2_HPO_4_, 1.47 mM KH_2_PO_4_), and lysed with 100 μl of Passive Lysis Buffer (Promega). Out of this lysate, 10 μl was used for the dual luciferase assay, and the rest was further lysed in an SDS-PAGE sampling buffer for the immunoblot analysis described below. The dual luciferase assay was conducted with Dual-Luciferase® Reporter Assay System (Promega) and a GloMax luminometer (Promega) according to the manufacturer’s protocols. The luciferase activity derived from the firefly luciferase reporter was normalized to that from the Renilla luciferase reporter.

### Immunoblot analysis

Immunoblot analysis was carried out as described previously (*23*). In short, proteins lysed in an SDS-PAGE sampling buffer were separated on a gel by SDS-PAGE, transferred to a Immobilon-P transfer membrane (Millipore), and probed with primary antibodies overnight at 4°C. The bound primary antibodies were detected by horseradish peroxidase-conjugated secondary antibodies in combination with an enhanced chemiluminescence detection system using Western Lightning Chemiluminescence Reagent (PerkinElmer Life Sciences) or ImmunoStar (Wako Pure Chemical Industries). Chemiluminescent images were acquired with ImageQuant LAS 4000 mini (GE Healthcare). The primary antibodies used were as follows: anti-FLAG antibody (F3165, Sigma, diluted to 0.8 μg/ml); anti-H3K4me3 antibody (07-473, Upstate, diluted 1:5000). The secondary antibodies used were as follows: horseradish peroxidase-conjugated anti-mouse IgG (074-1816, Kirkegaard & Perry Laboratories, diluted to 0.2 μg/ml) and horseradish peroxidase-conjugated anti-rabbit IgG (074-1516, Kirkegaard & Perry Laboratories, diluted to 0.2 μg/ml).

### Generation of transgenic zebrafish

To construct a Foxq2 transgene plasmid, the FLAG-Foxq2 coding sequence was amplified by PCR from the plasmid, drFoxq2/pCAG-FLAG (described in the section of Luciferase assay). The amplified fragment was then cloned with In-Fusion HD cloning kit into the pT2drCrx5.2kGP2ASix7 (*23*) digested by *Xho*I and *BamH*I. The resultant plasmid, named pT2drCrx5.2kGP2AFoxq2, was used for the generation of transgenic zebrafish. The *foxq2* transgenic zebrafish, ja78Tg, ja79Tg, and ja91Tg strains, were generated with the Tol2-based transgenesis system (*61*). The Tol2 transposase mRNA was transcribed from pCS-TP *in vitro* using the SP6 mMESSAGE mMACHINE Kit (Ambion) and purified with the RNeasy Mini kit (Qiagen). The purified Tol2 mRNA and the plasmid DNA were mixed and diluted to a final concentration of 25 ng/μl for each in 0.05% phenol red solution. About 1 nl of the DNA/RNA solution was injected into each of wild-type embryos at the one-cell stage. Fluorescence-positive embryos were isolated and raised to adulthood. The raised F0 founder fish were crossed with the wild-type fish, and subsequent F1 embryos were screened by the presence of fluorescence at four or five days post fertilization (dpf). The transgenic lines were established from individual F0 fish. The *foxq2-tg* larval fish and its siblings were genotyped by ocular EGFP fluorescence just before the sampling, and soaked in RNA*later* for RT-qPCR analysis or fixed with 4% PFA in D-PBS for cryosectioning. The GFP-negative siblings were used as a control. To unambiguously identify the transgenic larvae according to ocular EGFP expression, synthesis of melanin pigment was inhibited by treating embryos with the egg water containing 0.003% 1-phenol-2-thiourea (PTU) from 24 hours to 5 days after fertilization.

### Motif scanning

The *sws2* promoter (Fig. S4) was sequence-verified by traditional Sanger sequencing and used for the motif scanning. Fox and Crx binding profiles, each represented as a matrix consisting of nucleotide counts per position, *i.e.* position frequency matrix (PFM), were retrieved from the JASPAR CORE database (http://jaspar.genereg.net) (*62*). These retrieved binding profiles (total eleven profiles) were composed of one binding profile of Crx in mouse (*63*) and two binding profiles (FkhP and FkhS) for each of five Fox proteins in mouse (FoxA2, FoxL1, FoxK1, FoxJ1, and FoxJ3) (*64*). These frequency matrices were used to construct position-dependent letter-probability matrices that describe the probability of each possible letter at each position in the pattern with the simplest background model assuming that each letter appears equally frequently in the dataset. The *sws2* promoter was scanned for individual matches to each of the Crx and Fox motifs with FIMO v5.1.1 (*65*). Biased distribution of individual letters in the promoter sequences was normalized by a 0-order model of Markov background probabilities constructed with the fasta-get-Markov tool in the MEME suite v5.1.1 (*65*) using nucleotide sequences between 1000 bp upstream and 1000 bp downstream of a transcription start site for all protein-coding genes in zebrafish (genome assembly GRCz11, Ensembl Release 98). All motif occurrences with a P-value less than 1 × 10^−4^ are indicated in Fig. S4. The motif scanning results and matrix IDs of Fox and Crx in the database are included in Data S2.

### BLAST searches and phylogenetic analysis

For identifying *FOXQ2* genes in platypus and chicken, tBLASTn search (Ensembl web tools) was conducted against genome sequences of platypus (Reference Genome ID: mOrnAna1.p.v1) and chicken (Reference Genome ID: GRCg6a) using the amino acid sequence of forkhead domain of zebrafish Foxq2 (Ensembl protein ID: ENSDARP00000119225.2) as the query. We retrieved nucleotide sequences in intergenic regions showing the higher alignment score than any other regions encoding members of Fox families. The retrieved nucleotide sequences of *FOXQ2* genes were mapped onto two distinct genomic regions, but were adjacently located in the region of the same chromosome, where any other gene is not annotated. We thus assumed that these mapped regions are two exons of *FOXQ2* genes. Consistently, the forkhead domain of zebrafish Foxq2 is encoded in two exons. We then manually annotated exon-exon junctions of *FOXQ2* genes according to the GT/AG mRNA processing rule. The annotated cDNA sequence of the forkhead domain of *FOXQ2* gene was translated into a protein sequence and used for constructing a phylogenetic tree described in the next paragraph. The nucleotide sequences and annotations of *FOXQ2* genes are summarized in Data S3.

For constructing a phylogenetic tree, FOXQ2 amino acid sequences in spotted gar, medaka, coelacanth, and sparrow hawk were retrieved from Ensembl data base (Ensembl Release 101). FOXQ2 sequences in platypus and chicken identified by our blast searching were also used for the phylogenetic tree construction. Amino acid sequences of other members of Fox subfamilies were retrieved by performing tblastn searches (NCBI) against RefSeq RNA transcripts in purple sea urchin, zebrafish, chicken, platypus, and human using the amino acid sequence of forkhead domain of zebrafish Foxq2 as the query sequence with E-values < 1e−20. Among the retrieved cDNA sequences, members of representative Fox subfamily (FOXA, FOXB, FOXC, FOXF, FOXJ and FOXQ) were selected and translated into amino acid sequences. The resultant sequences of Fox proteins were aligned by multiple sequence alignment programs: G-INS-i program in MAFFT v7.471 under default settings (*66*). The aligned sequences were trimmed, remaining the sequences of the forkhead domain. Alignment gaps were manually inspected and deleted. The maximum likelihood tree was inferred by RAxML-NG v1.0.2 (*67*). The best tree was selected out of forty alternative runs on twenty random and twenty parsimony-based starting trees (--tree pars{20}, rand{20} option). The amino acid replacement models of Le-Gascuel (LG) with gamma distribution (G4) were selected using the Akaike information criterion implemented in ModelTest-NG version x.y.z (*68*). The bootstrap values were obtained from sampling 500 times. The amino acid sequences used for the construction of phylogenetic tree are listed in Data S3. Accession numbers for genome assemblies and Fox genes are also provided in Data S3.

### Statistical analysis

Sample sizes were determined based on prior literature and best practices in the field, and no statistical methods were used to predetermine sample size. A two-tailed unpaired *t*-test was used to determine the statistical significance between two datasets (Excel). Tukey–Kramer honestly significant difference test was used to determine the statistical significance among multiple datasets (the ‘stats’ package in R, version 3.6.1).

## Acknowledgements

We are grateful to members of the Fukada lab for valuable discussion, especially to M. Nagata for technical assistance. We thank the National BioResource Project (NBRP) Zebrafish for providing the transgenic line, and Dr. Shoji Kawamura, the University of Tokyo, for the zebrafish opsin transgenic lines, *Tg(−5.5opn1sw1:EGFP)*^kj9^, *Tg(−3.5opn1sw2:EGFP)*^kj11^, *Tg(RH2-2/GFP-PAC)*^kj4^ and *Tg(−0.6opn1lw1-lws2:GFP)*^kj19^. We also thank Dr. Takahiko Matsuda for providing the pCAG empty vector, Dr. Masato Kinoshita for the pCS2+hSpCas9 vector, and Dr. Wenbiao Chen for the pCS2-nCas9n vector. We also thank members of the FACS core laboratory, the University of Tokyo, for helping sorting cells. This work was supported in part by JSPS KAKENHI Grant Numbers JP16J01681 (to Y.O.), JP19K16196 (to T.S.), JP19K06758 (to D.K.) and JP17H06096 (to Y.F.) and also by a research grant from Research Foundation for Opto-Science and Technology (to D.K.). Y.O., T.S., D.K., and Y.F. conceived and designed the research. Y.O. and T.S. constructed plasmids and generated zebrafish lines. Y.O conducted all the other molecular lab work and computational analysis. D.K. and Y.F. supervised the project. Y.O., D.K., and Y.F. wrote the manuscript, and T.S. contributed to preparation of the draft. All authors approved the manuscript for publication. The authors declare that they have no competing interests. All data needed to evaluate the conclusions in the paper are present in the paper and/or the Supplementary Materials.

